# Structural mechanism for Bruton’s tyrosine kinase activation at the cell membrane

**DOI:** 10.1101/304097

**Authors:** Qi Wang, Yakov Pechersky, Shiori Sagawa, Albert C. Pan, David E. Shaw

## Abstract

Bruton’s tyrosine kinase (Btk) is critical for B-cell proliferation and activation, and the development of Btk inhibitors is a vigorously pursued strategy for the treatment of various B-cell malignancies. A detailed mechanistic understanding of Btk activation has, however, been lacking. Here, inspired by a previous suggestion that Btk activation might depend on dimerization of its lipid-binding PH-TH module on the cell membrane, we performed long-timescale molecular dynamics simulations of membrane-bound PH-TH modules and observed that they dimerized into a single predominant conformation. We found that the phospholipid PIP3 stabilized the dimer allosterically by binding at multiple sites, and that the effects of PH-TH mutations on dimer stability were consistent with their known effects on Btk activity. Taken together, our simulation results strongly suggest that PIP3-mediated dimerization of Btk at the cell membrane is a critical step in Btk activation.

---

Bruton’s tyrosine kinase (Btk), a Tec-family tyrosine kinase, is a peripheral membrane-binding protein present in all blood cells except for T cells and natural killer cells^1,2^. Btk participates in a number of receptor-mediated signaling pathways^2,6^. In B cells, where Btk’s biological functions are best understood, Btk activates phospholipase C-γ2, which produces second messengers that are essential for B-cell activation and proliferation. The upregulation of Btk is associated with various B-cell malignancies, including chronic lymphocytic leukemia and mantle cell lymphoma. Btk downregulation causes X-linked agammaglobulinemia (XLA), a severe disease of primary immunodeficiency^7^. Btk is also implicated in maintaining a number of autoimmune diseases, such as rheumatoid arthritis, systemic lupus erythematosus, and multiple sclerosis^8,9^. A structural characterization of Btk’s activation mechanism would significantly advance our understanding of Btk’s role in cell signaling and could facilitate the development of Btk-specific inhibitors,^10^ but such a characterization has thus far remained elusive.

Btk is autoinhibited in the cytoplasm, where it adopts a compact conformation similar to the inactive forms of the kinases c-Src and c-Abl,^11,12–14^ in which the SH2 and SH3 domains stabilize an inactive conformation of the kinase domain. In the case of Btk, its lipid-interaction module (the PH-TH module, composed of a pleckstrin homology (PH) domain and a Tec homology (TH) domain) acts in conjunction with the SH2 and SH3 domains to stabilize the inactive conformation of the Btk kinase domain.^11^ In B cells it has been shown that Btk activation is primarily regulated by phosphorylation upon membrane recruitment,^15,16^ which occurs through interactions between the canonical lipid-binding site of the PH-TH module and PIP_3_ membrane lipids, but the structural mechanism by which membrane recruitment leads to Btk activation is unknown.

Saraste and Hyvönen, after identifying a crystallographic dimer of the PH-TH module (often referred to as the “Saraste dimer”), suggested that dimerization of the PH-TH modules of membrane-bound Btk molecules may promote the trans-autophosphorylation of their kinase domains, leading to activation^17^. Dimers of the PH-TH module have never been directly detected in solution by biophysical methods, however, making their functional relevance uncertain. Membrane binding might be critical for PH-TH dimerization due to the increased local Btk concentration,^18,19^ but it is difficult to obtain atomic structural information about the protein on the membrane by traditional experimental means. Molecular dynamics (MD) simulations have enabled the observation of spontaneous binding events between proteins,^20–26^ providing an alternative, computational route for obtaining atomic structural information about protein oligomerization on membranes.

In this study, we use long-timescale MD simulations to examine whether and, if so, how the PH-TH module of Btk dimerizes on a membrane. In our simulations, which started from crystal structures of individual PH-TH modules and used no other prior structural information, the PH-TH modules spontaneously dimerized on the membrane into a single predominant conformation closely resembling the Saraste dimer. We observed that PIP_3_ allosterically stabilized the dimer conformation, binding not only at the canonical PIP_3_-binding site but at the peripheral IP_6_-binding site, which has been shown to be important for Btk activation in solution.^11^ At higher PIP_3_ concentrations, an additional PIP_3_ molecule bound at a site between the two PH-TH modules, interacting with both modules simultaneously, and there is evidence that this further stabilized the interface.

Our simulations also provide an explanation for the effects of multiple mutations in the PH-TH module that are known to be associated with Btk dysfunction: In our simulations, mutations that lead to downregulation of Btk and cause immunodeficiency diseases destabilized the PH-TH dimer, and a mutation that leads to the upregulation of Btk and causes cell overproliferation stabilized the dimer interface. These observations provide compelling evidence that the Saraste dimer is important for the regulation of Btk. Taking this together with our simulation results showing that PIP_3_ can allosterically mediate dimer formation, we further propose that PIP_3_ plays a physiological role as an allosteric activator of Btk by stimulating dimerization of the PH-TH modules and thus promoting trans-autophosphorylation of the kinase domains.

## Results

### The PH-TH module bound multiple PIP*3* lipids on the membrane

In order to establish the utility of our MD simulations for studying the behavior of the PH-TH module, we first showed that our simulations could reproduce essential experimentally observed interactions between the PH-TH module and soluble inositol phosphates (SI Text T1 and T2). We then studied the binding of the PH-TH module of Btk onto a membrane containing 94% POPC and 6% PIP_3_. We started the simulations by placing a PH-TH module in solution, about 10 Å away from the membrane surface, and positioned the module so that the canonical PIP_3_-binding site did not face the PIP_3_-containing leaflet of the membrane (Figure 1A).

**Figure 1.**
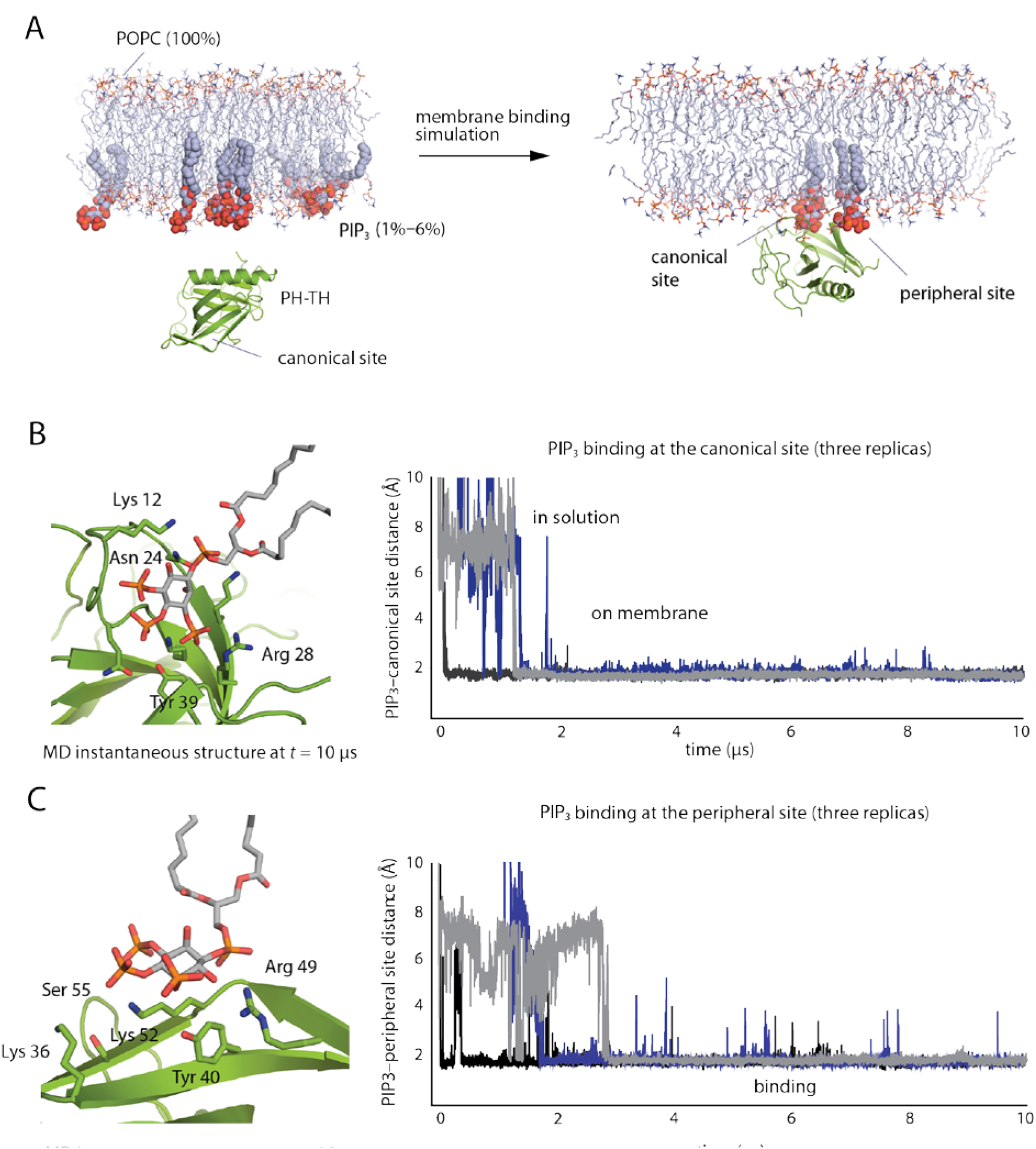
Spontaneous binding of the PH-TH module to the membrane. A. Cartoon illustration of a membrane-binding simulation setup (see SI Methods for details). B. Right panel: minimum distance between atoms in PIP_3_ lipids and in the canonical binding site for three independent membrane binding simulations. Left panel: a structure taken from one of these simulations (*t* = 10 μs) showing a PIP_3_ molecule bound at the canonical site. C. Right panel: minimum distance between atoms in PIP_3_ lipids and in the peripheral binding site for three independent membrane-binding simulations. Left panel: a structure taken from one of these simulations (*t* = 10 μs) showing a PIP_3_ molecule bound at the peripheral site.

Within 2 μs of simulated time, the PH-TH module was recruited onto the membrane, with a PIP_3_ molecule bound to the canonical site (Figure 1B). The binding pose of PIP_3_ at the canonical site resembles that of IP_4_ bound to the PH-TH module, as seen in crystal structures.^27^ Residues Asn 24, Tyr 39, Arg 28, and Lys 12 interacted with phosphate groups on the 2, 3, and 4 positions of the myo-inositol ring. In addition, Ser 21, Ser 14, and Lys 18 were also in close contact with the PIP_3_ molecule and formed transient interactions with PIP_3_ in the simulations. The bound PIP_3_ stayed at the canonical binding site until the end of simulations.

Once the PH-TH module was recruited onto the membrane through PIP_3_ binding to the canonical site, the peripheral IP_6_-binding site of the PH-TH module repeatedly bound a second PIP_3_ lipid (Figure 1C). Multiple PIP_3_ binding and unbinding events were observed at the peripheral site in individual trajectories, and the overall occupancy rate of the peripheral site was ~80% at 1.5% PIP_3_ concentration. The less-stable binding of PIP_3_ in the peripheral site is to be expected, since the peripheral site is relatively flat, whereas the canonical site has a well-defined groove where PIP_3_ binds. PIP_3_ assumed multiple binding poses at the peripheral site, with residues Lys 52, Tyr 40, and Arg 49 most frequently coordinating PIP_3_ binding; these are also the key residues experimentally observed to coordinate IP_6_ binding^11^ (Figure 1C). Residues Lys 36 and Ser 55, which are outside the area known to bind IP_6_, also interacted with PIP_3_ at the peripheral site.

Although MD simulation studies have shown that several PH domains bind anionic lipids at non-canonical binding sites^28—31^, the peripheral site identified in our study has not been previously reported as a lipid-binding site for Btk. In our simulations we also observed PIP_3_ repeatedly occupying a third binding site, located on the other side of the IP_4_-binding loop (Figure S1). We have not studied PIP_3_ binding in this pocket in detail, however, due to its low occupancy rate (~30%) in our simulations.

Our simulations do not rule out the possibility that other types of anionic lipids, such as PIP_2_ or PS, might also bind at these regions of the PH-TH module. In this work, we chose to exclusively study PIP_3_ due to the critical signaling role it is known to play in mediating B-cell activation. The finding that the PH-TH module can simultaneously bind multiple PIP_3_ lipids on the membrane has, to our knowledge, not been noted in prior studies, and has important implications for Btk activation on membranes, as we will show in the results below.

### The PH-TH modules spontaneously dimerized on the membrane with a single predominant interface

PIP_3_-bound PH-TH modules diffused freely on membranes in our simulations, and we were thus able to study the encounter process of two membrane-bound PH-TH modules. Here we show that PH-TH modules can indeed spontaneously dimerize on membranes, with a single dimer interface predominating.

We began by performing conventional MD simulations of two separate, non-membrane-bound PH-TH modules and a piece of membrane containing 6% PIP_3_ (Figure 2A). In 20 independent 20-μs trajectories, the individual modules always bound with the membrane, and a variety of PH-TH dimers subsequently formed, in each case within the first few microseconds of simulation time (Figure S2). Some dimer interfaces, despite having small interface areas (~500 Å^2^) and lacking hydrophobic packing interactions and hydrogen bonds, stayed bound through the end of the 20-μs simulations (Figure S2). The fact that such dimer interfaces did not dissociate in the simulations is not unexpected: Observing dissociation of protein-protein complexes is a well-known challenge for conventional MD simulations, since experimentally observed dissociation events often occur on a longer timescale than is accessible to MD simulations.^26^

**Figure 2.**
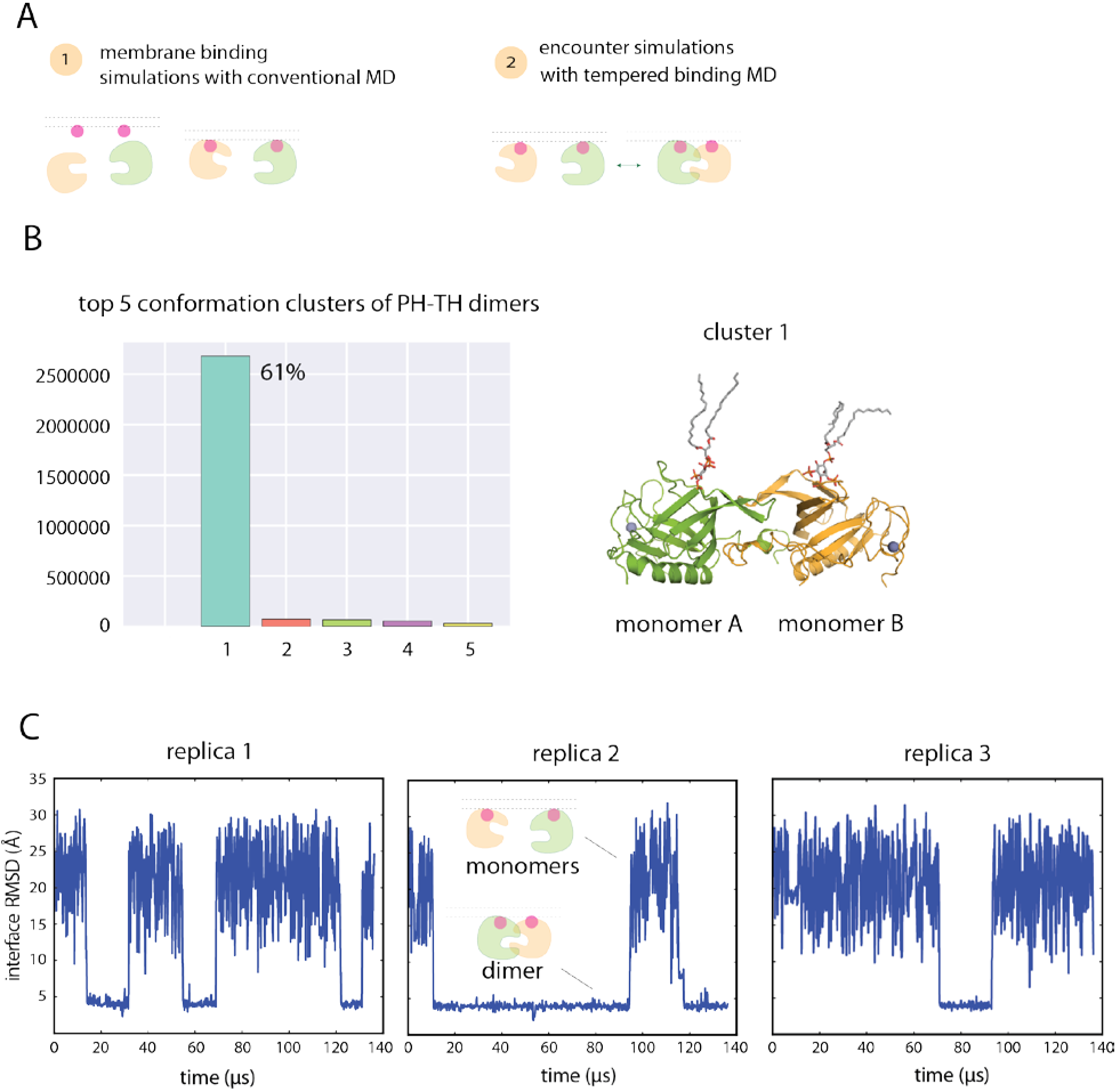
Dimerization of the PH-TH module on the membrane. A. Cartoon illustration of an encounter simulation setup (see SI Methods for details). B. Conformation clusters of PH-TH dimers formed in the simulations (see SI Methods for details). A representative structure of the most populated dimer conformation is shown on the right. (The PIP3 lipids bound in the canonical binding sites are also shown.) C. Reversible dimerization of two membrane-bound PH-TH modules on a membrane that contains 1.5% PIP_3_ and 98.5% POPC in tempered binding simulations. The RMSD at the Saraste interface is calculated for all interface residues (residues 11, 42, 44, 92, and 95) of the two modules with respect to the Saraste dimer interface conformation seen in a crystal structure of the PH-TH module (PDB ID: 1BTK). (This same interface RMSD calculation is used for the RMSD plots in all other figures unless stated otherwise.)

In order to sample PH-TH module conformations more extensively than would be possible using conventional MD simulations of reasonably achievable length, we employed an enhanced sampling method called “tempered binding.”^26^ This technique intermittently weakens the interactions between the two proteins (and thus all potential PH-TH dimer interfaces), thus increasing the frequency of dimer dissociation and accelerating the exploration of different protein-protein interfaces. The method nonetheless rigorously preserves the property of Boltzmann sampling at each interaction strength, with unscaled portions of the trajectory sampled from the same distribution as a conventional MD simulation with an unmodified Hamiltonian (see SI Methods for details).

We performed 24 tempered binding simulations, each based on three replicas, with a total of ~6 ms of simulated time for all simulations combined. All tempered binding simulations started from the same configuration—in which two PH-TH modules were bound to the membrane but were not in contact with each other—but varied in membrane PIP_3_ concentration, tempering strength, and protein-backbone-correction strength.

In these tempered binding simulations, the two PH-TH modules diffused freely on the membrane, and multiple association/dissociation events between the two modules were observed. We then analyzed all dimer conformations assumed by the PH-TH modules in these simulations: The two modules were in contact in 4 million of the 6 million total frames extracted from the 24 simulations, and we began by clustering the instantaneous structures in these 4 million frames according to their structural similarity (Figures 2B and S3). The largest cluster of these PH-TH dimer structures contained 61% of total dimer structures, while the next-largest cluster contained less than 1%; given the small size of all but the largest, all further structural analysis was based only on the most populated cluster (Figure 2B). Notably, clustering only those portions of the trajectory in which the protein-protein interactions were untempered led to the same predominant cluster, supporting the relevance of this dimer cluster under untempered conditions.

Dimers in the predominant cluster formed repeatedly in all 24 simulations (across all tempering parameters and PIP3 concentrations). The dissociation time for these dimers varied from simulation to simulation, ranging from at least 15 μs of simulation time to up to 500 μs, which was the length of the longest simulations. We also observed multiple reversible dimerization events for these dimers in individual trajectories under certain tempering conditions (Figure 2C).

Dimer structures in other clusters, on the other hand, were short-lived, and most of them dissociated within a few microseconds of simulation time.

### The predominant PH-TH dimer interface resembles the Saraste dimer interface

The most populated PH-TH dimer interface in our simulations resembles the Saraste dimer observed in crystal structures (Figure 2B): The root-mean-square deviations (RMSDs) of the PH-TH dimer interfaces in the largest cluster range from 2.5 to 4.5 Å compared to the Saraste dimer. The Saraste-like dimer interfaces that formed in our simulations had two distinct conformations (Figure 3A and 3B), with the primary structural difference between them occurring at the ß3-ß4 hairpin (Figure 3B). In the conformation closest to the Saraste crystal structure, with an RMSD at the dimer interface of only ~2.5 Å (Figure 3A), the interface is tightly packed. In the other conformation, the dimer interface is loosely packed, and the interface regions of the individual PH-TH modules are partially closed (Figure 3B): The Phe 44 of each module remains packed against its own module’s Ile 92 and Ile 95 residues, as is also the case in simulations of a single PH-TH module. The RMSD of the second conformation at the dimer interface is ~4.0 Å compared to the Saraste dimer crystal structure.

**Figure 3.**
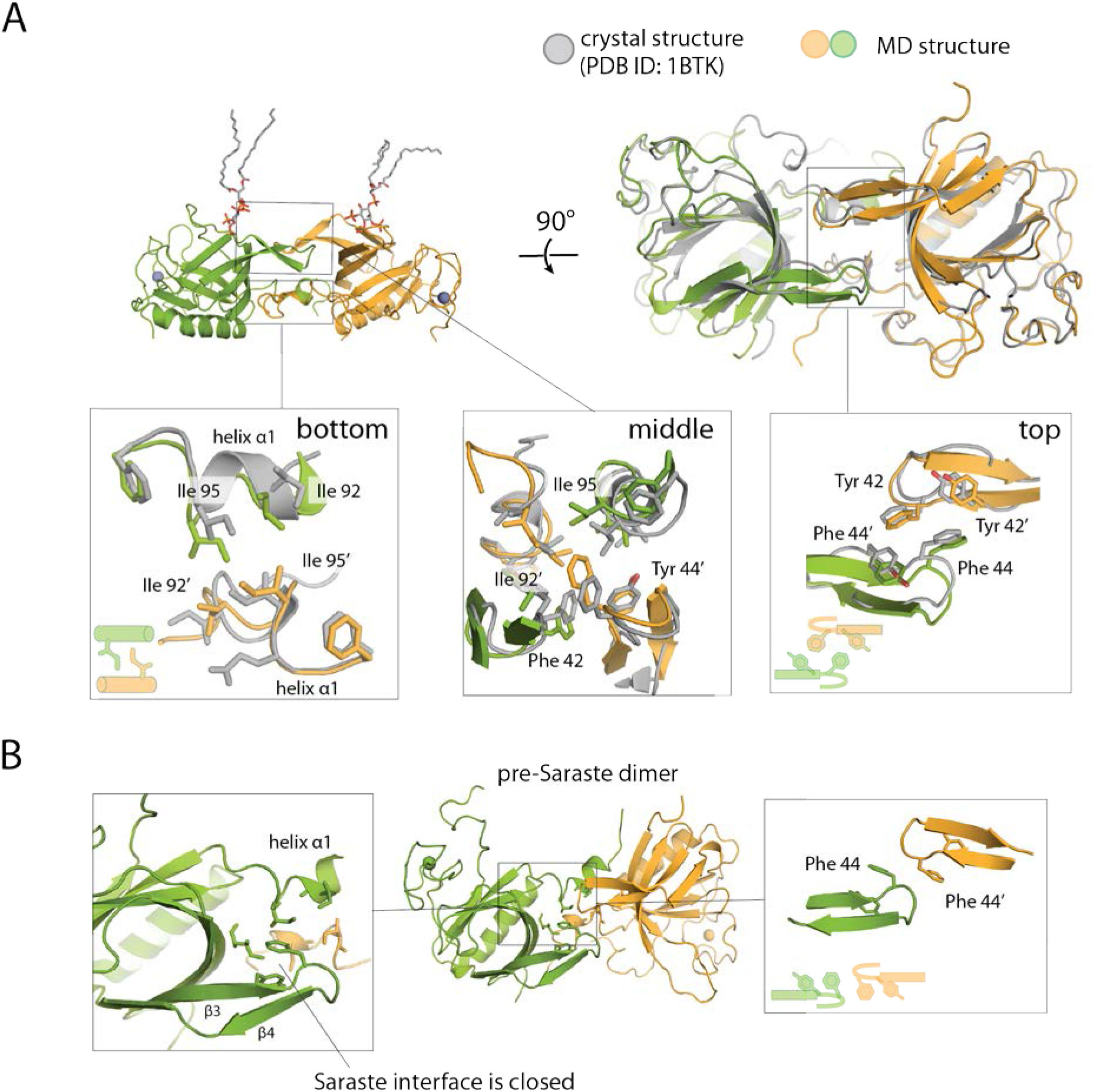
Structural analysis of Saraste dimers formed on membranes. A. Structural comparison of Saraste dimers formed in the encounter simulations and in the Saraste dimer crystal structure (PDB ID: 1BTK). B. An instantaneous structure of the pre-Saraste interface formed in the encounter simulations.

This loosely packed conformation of the Saraste interface has not been observed in crystal structures of the PH-TH dimer. Nevertheless, we find this conformation very informative: It provides evidence for a structural mechanism by which the tightly packed Saraste dimer interface can form while the interface regions of the individual PH-TH modules remain partially closed (SI Text T1). In our simulations of an individual PH-TH module, residues at the Saraste interface region rarely adopted the open conformation, which is the conformation compatible with directly forming the tightly packed Saraste interface with another module. We thus did not expect the direct formation of the Saraste dimer interface to occur frequently in our simulations, since it would require the spontaneous opening of the interface regions of both PH-TH modules simultaneously during the encounter process. Indeed, none of the tightly packed Saraste dimer interfaces we observed in our tempered binding simulations formed through the direct encounter of the two modules with open Saraste-interface regions: All Saraste dimer interfaces formed by way of conformational transitions through the loosely packed interface. We thus call the loosely packed interface the “pre-Saraste” interface. Furthermore, in one of our conventional MD simulations of two initially separated PH-TH modules on a membrane, we also observed the PH-TH modules spontaneously form the pre-Saraste dimer, strongly suggesting that the pre-Saraste dimer is an obligatory intermediate state between the unbound modules and the tightly packed Saraste dimer.

### The stability of the Saraste interface was sensitive to mutations on the PH-TH module

The observation in our simulations that the PH-TH modules spontaneously dimerized with a single predominant interface on the membrane, where Btk performs its biological functions, suggests that this Saraste dimer interface may be biologically relevant. Additionally, in solution the transient dimerization of PH-TH modules at the Saraste interface in the presence of IP_6_ can activate Btk molecules by promoting trans-autophosphorylation of their kinase domains, and mutations that destabilize the Saraste interface slow Btk activation in solution.^11^ Together, these findings suggest that the stability of the Saraste dimer may regulate activation of membrane-bound Btk. To test this hypothesis, we used MD simulations to study two mutations on the PH-TH module that are known to affect Btk activity, but for which a structural explanation is lacking. In our simulations, we examined whether and, if so, how these mutations influence the stability of the Saraste dimer on a membrane.

Initially we explored using tempered binding simulations to study the thermodynamic equilibrium between a Saraste dimer and the unbound modules, but the results were inconclusive due to the small number of association and dissociation events that could be observed on our simulation timescale. Instead, we proceeded by qualitatively analyzing the potential structural consequences for Saraste dimer stability when different mutations were introduced. In this set of simulations, we used the tightly packed conformation of the Saraste dimer on a membrane as a starting structure and performed tempered binding simulations until the dimer dissociated (Figure 4A).

**Figure 4.**
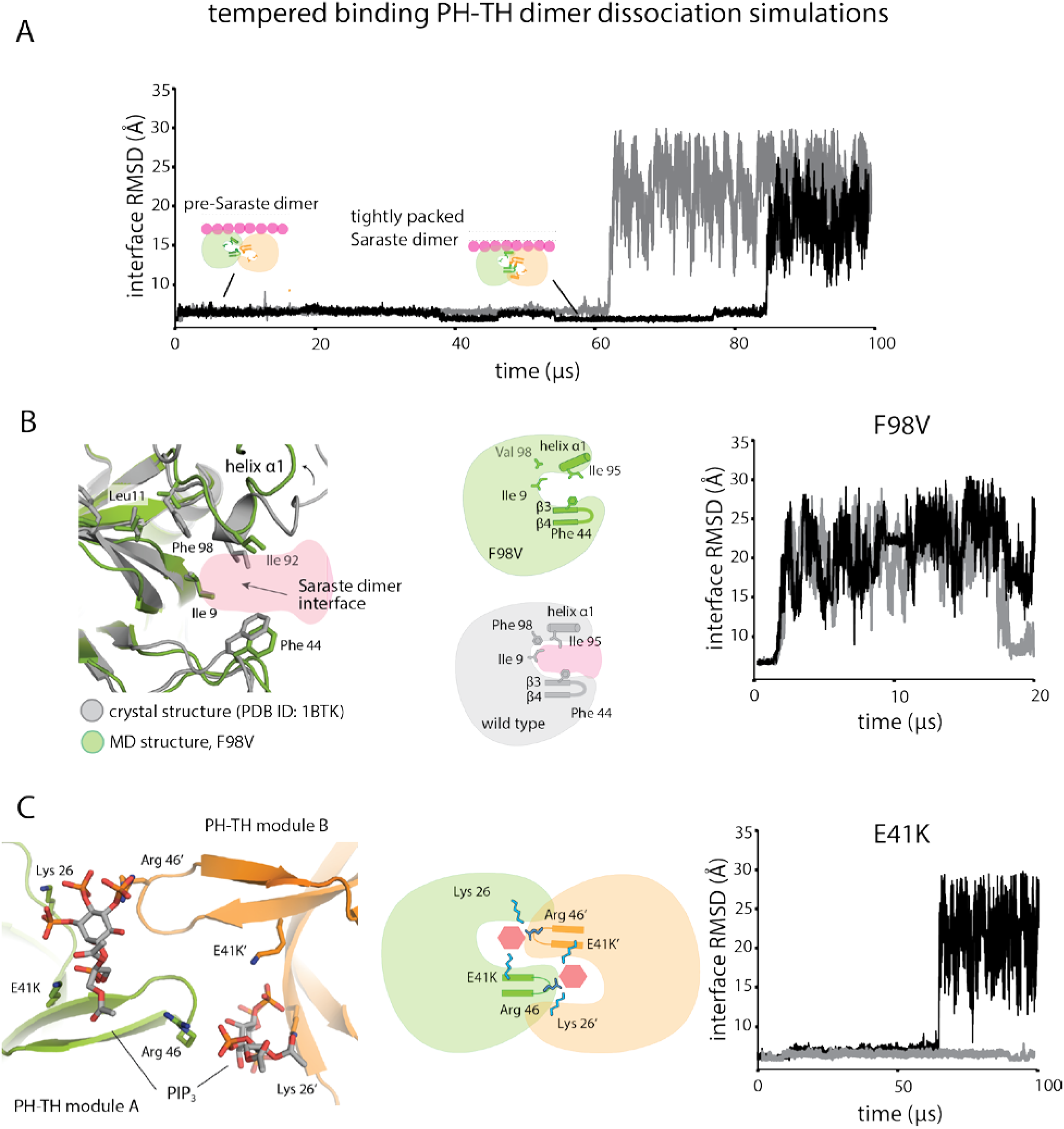
Stability analysis of Saraste dimer variants on a membrane. A. Representative tempered binding simulation trajectories showing dissociation of the tightly packed Saraste conformation in two individual trajectories. The tempered binding simulations started from a Saraste dimer on a membrane containing 6% PIP3 and 94% POPC. B. Stability analysis of the F98V mutant, starting from the Saraste dimer conformation, on a membrane. An instantaneous structure from a simulation (*t* = 20 μs) shows the local conformational rearrangement that occurred at the Saraste interface before the dimer dissociated. Representative tempered binding simulation trajectories show the dissociation of the F98V dimer in two individual trajectories. C. Stability analysis of the E41K mutant, starting from the Saraste dimer conformation, on a membrane. An instantaneous structure from a simulation (*t* = 100 μs) shows PIP3 bound to residue Lys 41 at the bridging site before the dimer dissociated. Representative tempered binding simulation trajectories show the dissociation of the E41K dimer in two individual trajectories.

We first studied the loss-of-function mutant F98V, which is located in the PH domain and has been identified in patients with the immunodeficiency disease XLA.^32^ The molecular basis for the loss of function produced by F98V is not clear: Phe 98 is involved neither in the hydrophobic-core packing interactions of the PH-TH module nor in directly mediating PIP_3_-binding interactions, suggesting that the loss-of-function effect of F98V is unlikely to be related to misfolding of the individual PH-TH module or to interference with its membrane recruitment. One effect we did observe in our simulations, however, was that substituting Phe with the less-bulky Val loosened the packing between residues Leu 11 and Leu 9 (Figure 4B). Leu 9 is part of the Saraste interface, suggesting that the loosened packing at the end of the α1 helix, allosterically produced by F98V, could affect the stability of the Saraste interface. Consistent with this notion, in our simulations we observed that the F98V dimer was not stable in the tightly packed Saraste interface conformation, quickly relaxed into the pre-Saraste conformation, and then dissociated into separate PH-TH modules (Figures 4B and S4). In addition to F98V, we observed that the Saraste interface was less stable in another mutant known to cause XLA, F25S (Figure S4).

We next studied the effects of the mutation E41K, a gain-of-function mutation in the PH-TH module of Btk^33^. Glu 41 is located on the ß3—ß4 hairpin, near the Saraste interface, and its mutation to lysine constitutively activates Btk and causes cell overproliferationAlthough previous studies have shown that the membrane localization of the E41K mutant is more significant than that of the wild-type protein^34–37^, the structural basis of this effect, and the interactions between E41K dimers and membranes after the mutant is recruited onto the membrane, are not well understood. In our simulations, we observed Lys 41 repeatedly bind a third PIP_3_ lipid at the top of the ß3-ß4 hairpin, confirming the notion, based on previous crystallographic studies, that the E41K mutant can bind an additional PIP3 molecule^27^. Although the stability of the E41K dimer was qualitatively comparable to that of the wild-type dimer in our simulations, the PIP_3_ that bound at the top of the ß3-ß4 hairpin interacted simultaneously with both modules in the Saraste dimer (leading us to term this binding location the *bridging site)*, revealing a structural mechanism that could further strengthen Saraste-interface stability (Figure 4C). We thus speculate that the activating effect of E41K could be the result of additional PIP3 molecules stabilizing the Saraste interface once the dimer has formed on the membrane, which could further promote the trans-autophosphorylation of Btk kinase domains.

Finally, we note that, as expected, Saraste dimers with mutations at the dimer interface (I9R/L29R, Y42R/F44R) either barely stayed in the tightly packed Saraste interface conformation or dissociated immediately in our simulations (Figure S4). These mutants have been previously shown to impair Btk activation by IP_6_ in solution,^11^ and our simulation results strongly suggest that they are also likely to impede Btk activation on membranes.

The qualitative correlation between the stability of the mutated Saraste dimers and the functional consequences of these mutations suggests that the Saraste interface plays an important role in regulating Btk activation.

### The stability of the Saraste dimer interface was also sensitive to the amount of PIP_*3*_ in the membrane

In B cells, downregulation of Btk activity causes immunodeficiency diseases, and upregulation of Btk activity is correlated with autoimmune diseases and various blood cancers, suggesting that the activation of Btk must be tightly controlled to maintain normal B-cell functions. The observation that Btk function is very sensitive to the stability of the Saraste dimer interface suggests that the stability of the Saraste dimer itself might be tightly controlled in cells. We now describe a set of simulation observations that collectively suggest that PIP_3_, the anchor of Btk on membranes, is a potential allosteric regulator for Btk that may control the stability of the Saraste interface.

The first insight came from analyzing the tempered binding simulations discussed above, in which we found that Saraste dimers that formed on membranes with 6% PIP_3_ lipid content did not dissociate as frequently as those that formed on membranes with 1.5% PIP_3_ (as seen in Figures 5A and 2C, respectively). Although we were unable to quantitatively compare the dissociation constants of the Saraste dimer at 1.5% and 6.0% PIP_3_ membrane lipid concentrations due to the small number of association and dissociation events, we found that fluctuations at the pre-Saraste interface, which preceded dissociation in our simulations, were more pronounced in simulations with 1.5% membrane PIP_3_ content than in those with 6% PIP_3_ (Figure 5B). Dimer conformations with RMSDs greater than 4.5 Å from the Saraste dimer crystal structure at the interface were repeatedly visited in the 1.5% PIP_3_ condition (Figure 5B). These conformations resembled the partially broken Saraste dimer conformations seen in our solution-based conventional simulations (SI Text T3 and Figure S5). The large, relatively frequent conformational fluctuations at the dimer interface that repeatedly visit a structure on the dissociation pathway suggest that the dimers were weaker in the 1.5% PIP_3_ condition. In simulations with 6% membrane PIP_3_ content, the RMSD from the crystal structure at the Saraste interface consistently stayed below 4.5 Å. Overall, our observations provide strong evidence that the Saraste dimer was more stable in the 6.0% PIP_3_ lipid condition than in the 1.5% PIP_3_ lipid condition.

**Figure 5.**
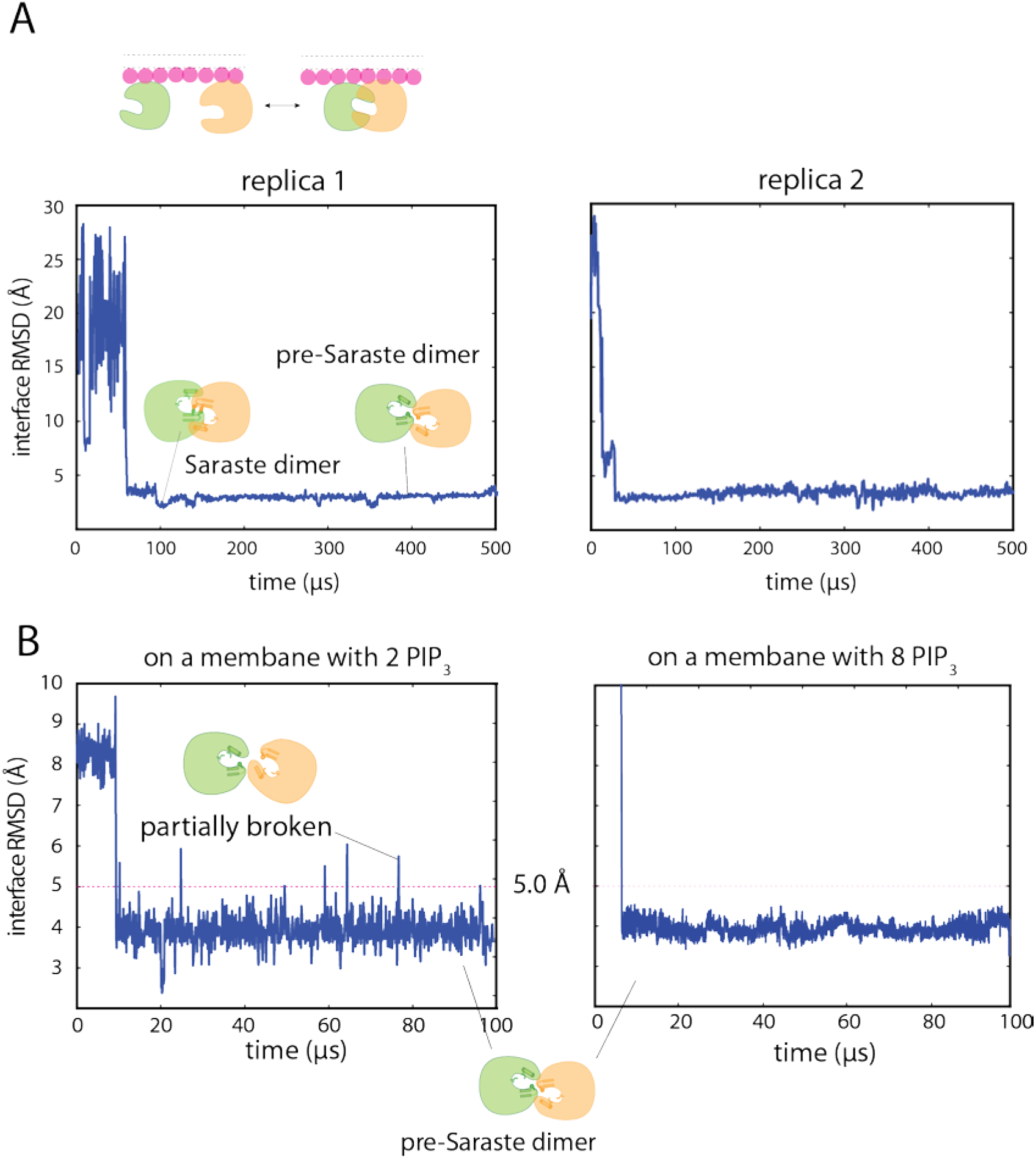
Fluctuations at the pre-Saraste dimer interface. A. Representative tempered binding simulation trajectories of two PH-TH modules on a membrane containing eight PIP_3_ lipids. B. Fluctuations at the pre-Saraste dimer interface that formed on membranes with two and eight PIP3 lipids (1.5% and 6% membrane PIP3 content, respectively).

### PIP_3_ binding at the canonical and peripheral sites of the PH-TH modules reduced fluctuations at the Saraste dimer interface

Some polyphosphoinositides, such as PIP_2_ and PIP_3_, are known to spontaneously form clusters under certain conditions^38^. One possible explanation for the apparently rare dissociation of the Saraste dimer in the 6% PIP_3_ condition is that PIP_3_ clusters spatially constrained the PH-TH dimer on the membrane, leading to infrequent dimer dissociation. We examined the localization pattern of PIP_3_ in our simulations to investigate this possibility, but found that no stable PIP_3_ clusters formed in any of our simulations. Indeed, the PIP_3_ molecules were quite isolated from each other, as a result of binding to distinct sites on the PH-TH module. We found that this binding of PIP_3_ at both the canonical site and the peripheral site of PH-TH modules had profound effects on the stability of the Saraste interface, suggesting that PIP_3_ plays a role in regulating the PH-TH dimer allosterically.

To study more rigorously whether both the canonical and peripheral sites are involved in the stabilizing effect of PIP_3_ binding described in the previous section, we performed another set of tempered binding simulations, varying only the number of PIP_3_ lipids in the membrane (from one to eight), and analyzed the fluctuations at the dimer interface after the formation of the pre-Saraste interface. We first observed that the Saraste dimer interface was not stable in solution, and that PIP_3_ binding significantly reduced fluctuations at the Saraste interface (Figure S5A). PIP_3_’s stabilizing effect on the Saraste interface became evident when comparing the interface conformations when one or both of the two canonical sites in the Saraste dimer had bound PIP_3_ ligands (Figures 6 and S5A): In simulations with only one PIP3 molecule in the membrane, one PH-TH module in the Saraste dimer was attached to the membrane at the canonical site, while the other module remained in solution. We observed that conformations with large RMSDs (4.5 Å or more) from the Saraste crystal structure at the dimer interface appeared frequently in simulations with only one PIP3 molecule (Figure 6). When two PIP3 lipids were present, occupying both canonical binding sites in the Saraste dimer, the population of conformations with large RMSDs at the Saraste dimer interface shrank from 16% to 8%, and the population of conformations with small RMSDs (3.0 Å and below) at the Saraste dimer interface increased to 7% (Figure 6).

**Figure 6.**
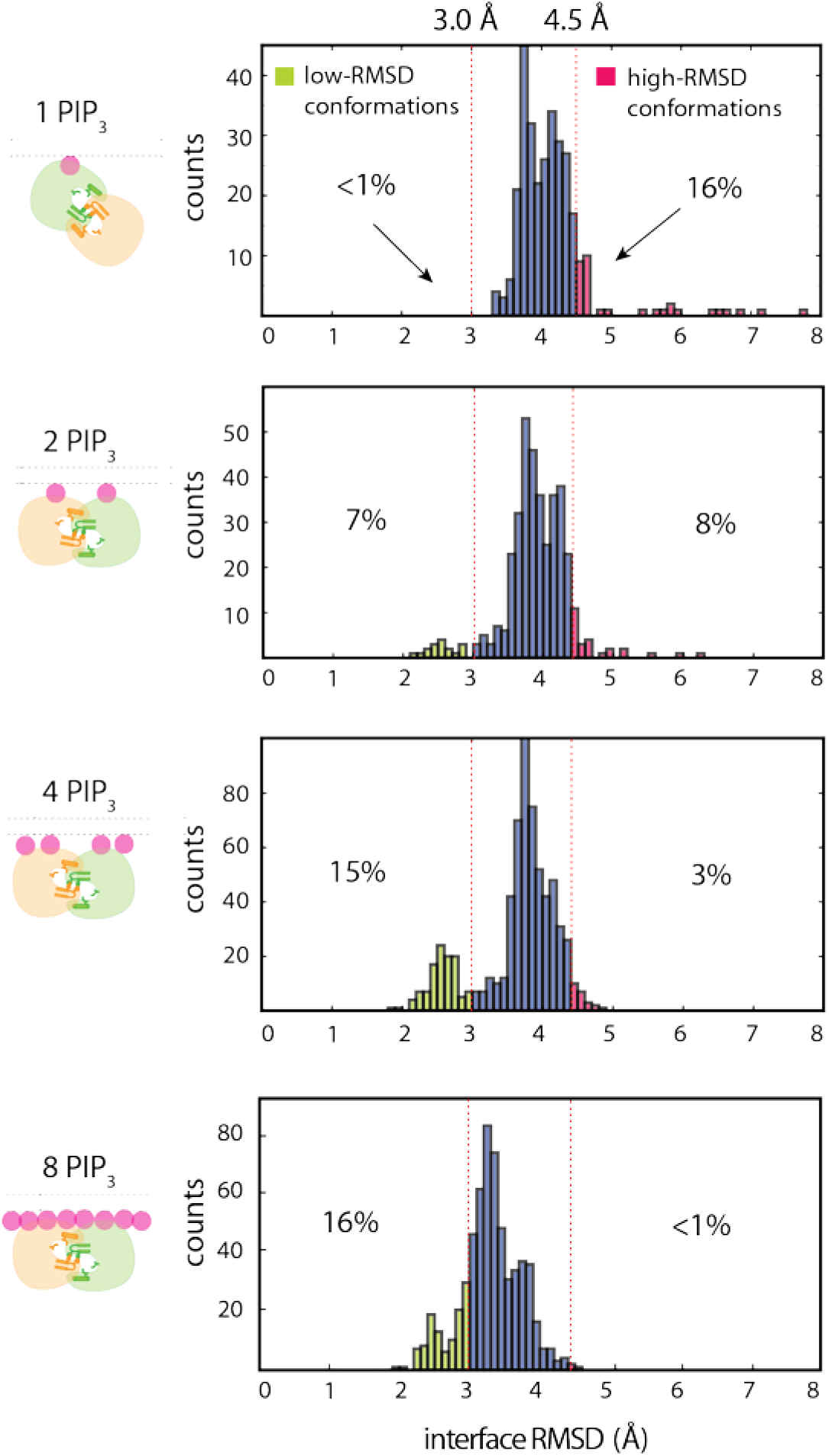
Sensitivity of fluctuations at the pre-Saraste dimer interface to membrane PIP_3_ concentration. Fluctuations at the pre-Saraste dimer interface are shown, measured under various membrane PIP_3_ concentrations. The histograms of the RMSD of the interface residues are calculated from 100-μs simulation trajectories initiated from Saraste dimers that originally formed in tempered binding simulations; only data from the portion of the trajectories where the interactions were unscaled is used.

We then observed that with four PIP_3_ lipids in the membrane, the stability of the Saraste dimer improved further (Figure 6). In this set of simulations, PIP_3_ lipids typically occupied both the canonical and peripheral sites in the Saraste dimer: The occupancy rate at each peripheral binding site was ~80%. In these simulations, the population of conformations with large RMSDs at the dimer interface fell further, to 3%. The population of conformations with small RMSDs at the Saraste dimer interface increased from 7% to 15%, consistent with the notion that binding of PIP_3_ at the peripheral site further stabilizes Saraste interface.

In the presence of membranes with eight PIP_3_ molecules (6% membrane PIP_3_ content), the canonical and peripheral sites of the Saraste dimer were almost continuously occupied in our simulations. We observed a further reduction in the population of conformations with large RMSDs, from 3% to <1%. In addition, we found that the peak RMSD distribution was shifted from 3.7 Å to 3.2 Å, suggesting that the PIP_3_ occupying the peripheral site of each PH-TH module stabilizes the Saraste interface.

Further evidence for the importance of the PH-TH module’s peripheral binding site in stabilizing the Saraste interface is seen in simulations we performed to reveal the thermodynamic equilibrium between the tightly packed Saraste conformation and the more loosely packed pre-Saraste conformation (Figure S6A; see SI Methods for details). In simulations with 6% PIP_3_ content, the PH-TH dimer reversibly converted between the pre-Saraste conformation (45%) and the tightly packed Saraste conformation (55%) before dissociating (Figure S6B). The free energy difference of the transition between the two Saraste conformations was 05 ± 0.2 kcal mol ^−1^. In the presence of a membrane with 1.5% PIP_3_ content, we found that the fraction of the tightly packed Saraste dimer fell to 40%, and the free energy difference between the two Saraste conformations decreased to −0.4 ± 0.15 kcal mol^-1^, consistent with the notion that the tightly packed Saraste dimer conformation is less stable in the presence of a membrane with 1.5% PIP_3_ content.

### PIP_3_ binding at the canonical site and the peripheral site rigidified the Saraste dimer interface in an individual PH-TH module

Our simulations point to important roles for the canonical site and the peripheral site in controlling Saraste dimer formation, but the precise mechanisms by which PIP_3_ binding strengthens Saraste dimer stability are not clear. After the binding of PIP_3_ at the two sites, we observed no obvious structural change in the PH-TH module that might lead to a more stable dimer conformation. We thus investigated the possibility that the binding of two PIP_3_ molecules could restrict the range of conformations that the Saraste interface region can adopt.

To investigate this possibility, we studied the conformational dynamics of an individual PH-TH module under various conditions: In solution-based simulations, the RMSD from the Saraste crystal structure at the dimer interface ranged from 2.5 Å to 6.5 Å, with a peak distribution at ~4.8 Å (Figure 7A). When the PH-TH module was recruited onto a membrane with only one PIP_3_ lipid occupying the canonical binding site, the fluctuations at the Saraste dimer interface decreased, with the distribution peak shifting from 4.8 Å to 4.2 Å RMSD (Figure 7A). In the presence of two PIP_3_ lipids, the peripheral site was also occupied, and the peak distribution decreased further, to 3.5 Å (Figure 7A). These observations indicate that the conformational dynamics of the PH-TH module are more restrained when both the canonical site and the peripheral site are occupied by PIP3.

**Figure 7.**
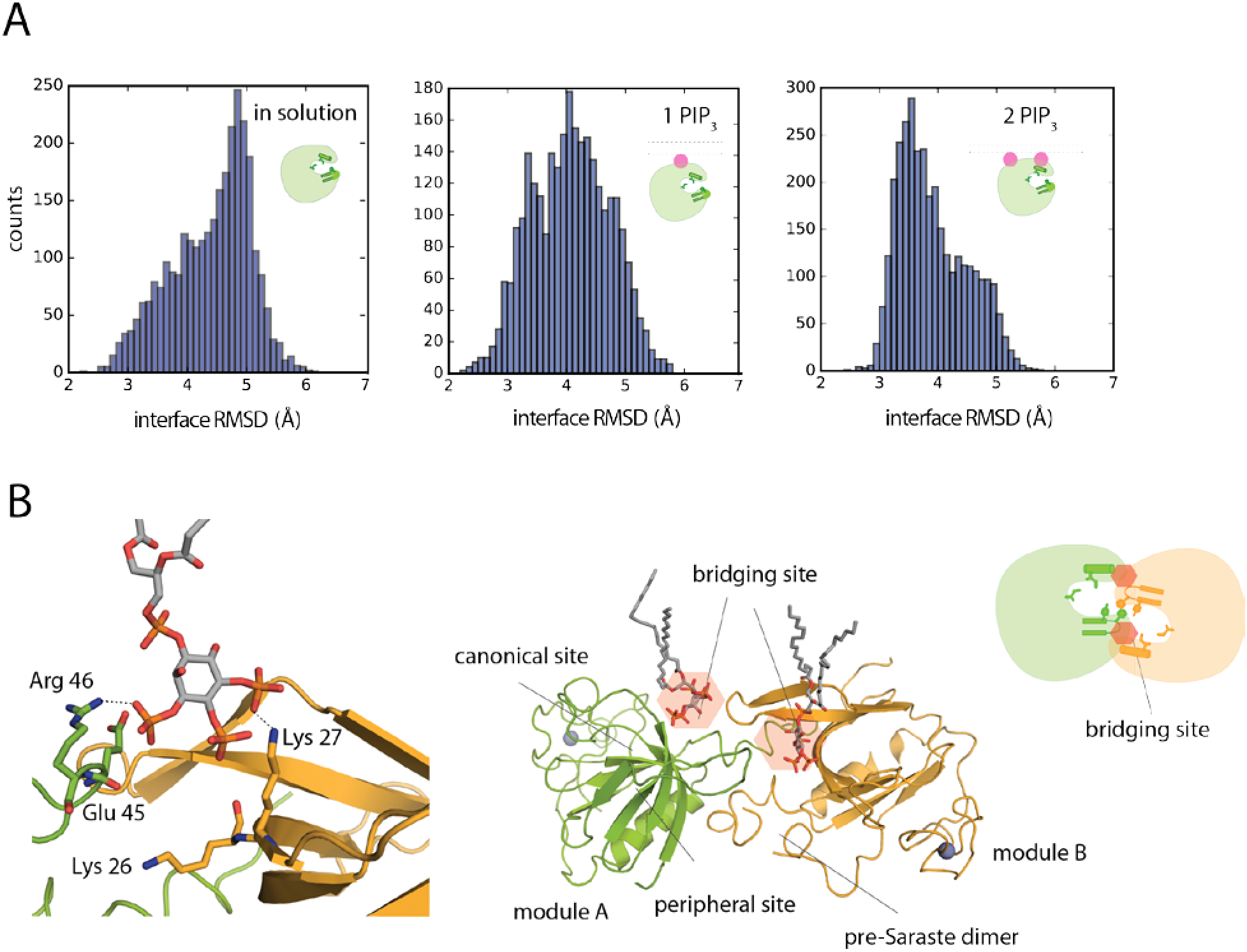
Speculation on the mechanisms by which binding multiple PIP_3_ molecules stabilizes the Saraste dimer on the membrane. A. Fluctuations at the Saraste dimer interface region in an individual PH-TH module in conventional MD simulations in solution and on the membrane. B. An instantaneous structure from a tempered binding simulation showing that a PIP3 lipid can simultaneously interact with both modules in the Saraste dimer at the bridging site, which formed near the dimer interface.

### At high concentrations, PIP_3_ bound at a bridging site, where it interacted with both PH-TH modules in a Saraste dimer

In the presence of membranes with 6% PIP_3_ content, the canonical and peripheral sites of the Saraste dimer were mostly occupied in our simulations. Notably, at this concentration we found that the area between the tip region of the ß3—ß4 hairpin of one PH-TH module and the edge of the canonical binding site of the other module also bound a PIP3 molecule (Figure 7B). In an earlier section, we suggested that PIP3 binding at this bridging site might explain the activating effect of the E41K mutant; in the presence of a membrane with 6% PIP_3_ content, the bridging site was transiently occupied by a PIP3 lipid even in the absence of this mutation. In one representative structure extracted from our simulations, the phosphates at positions 1 and 4 of the inositol ring of PIP_3_ interacted simultaneously with the Arg 46 residue of one module and Lys 27 of the other, which we label modules A and B, respectively (Figure 7B). The hydroxyl group at position 2 also occasionally formed a hydrogen bond with Glu 45 of module A. The average occupancy rate of the bridging site was low, at ~40% when the membrane had 6% PIP_3_ content, suggesting that PIP_3_ binding at this site is of low affinity. We speculate that the bridging site might contribute to Saraste dimer stability only in the presence of a large number of PIP_3_ lipids.

The two mechanisms we have described—stabilization by PIP_3_ of the individual PH-TH modules in the Saraste dimer—compatible conformation and, at high concentrations, the interaction of PIP_3_ with both modules simultaneously to stabilize the dimer once it has formed—together support the notion that PIP_3_ can allosterically stabilize the Saraste dimer. Little is known about the actual local concentration of PIP_3_ near B-cell receptors, where Btk primarily functions. PIP_2_, the precursor molecule of PIP_3_, has been found in protein microdomains with a local concentration as high as 82%^39^. PIP_3_ has also been found in protein microdomains, with a local concentration up to 16%.^40^ In our simulations, the concentration of PIP_3_ was 1%-6%, falling within the range of potential PIP_3_ concentration in cells. It is thus reasonable to expect that both mechanisms observed in our simulations may occur in nature.

## Discussion

### A structural model of Btk activation on membranes

Based on our atomic-level MD simulations, we have presented two important findings, which together provide a potential mechanism for Btk activation (Figure S7). First, we observed that Btk PH-TH modules spontaneously dimerized on membranes into a single predominant conformation, one which resembles the Saraste dimer. This finding provides a structural explanation for the effects of mutations in the PH-TH module known to cause severe Btk dysfunction, as these mutations affected the stability of the dimer.

The other major finding we have presented in this work is that PIP_3_ can allosterically stabilize the Saraste dimer interface, and may thus play a regulatory role in Btk activation. In prior models of Btk regulation, PIP_3_ has been considered a membrane anchor for the kinase, but no further regulatory role for PIP_3_ has previously been proposed. Our simulations provide evidence that PIP_3_ controls Saraste dimer stability, which may in turn have a profound impact on Btk activation: In order for Btk to be activated by trans-autophosphorylation, the two kinases need to come close together and form a transient enzyme-substrate complex. At a single-molecule level, the lifetime of the enzyme-substrate complex ought to be longer than the time required for transferring the phosphate from ATP to the substrate tyrosine. Increasing the lifetime of the Saraste dimer would thus make the trans-autophosphorylation reaction more efficient. (Although for this work we have not studied in detail whether PIP_3_ binding affects the on-rate of PH-TH dimer formation, which would also promote Btk activation, there is some evidence from our simulations that tends to support this proposition (SI Text T4 and T5).)

PH domains, which act as membrane-binding modules and protein-protein interaction modules, are frequently found in the human genome.^41,42^ Dimerization of membrane-bound PH domains has received little attention in the past, but PH domain dimerization may play an important role in enzyme regulation. Recently, the PH domain of phosphoinositide-dependent kinase-1 (PDK1), a serine/threonine kinase that is important for cell differentiation and proliferation pathways, was shown to dimerize on a supported bilayer system.^43^ The structural basis for this dimerization is not clear, but it has been speculated that the PDK1 dimer represents an inhibitory state of the kinase that is not capable of substrate binding.^44^ This notion, together with the activating role of the PH-TH module of Btk proposed in this paper, leads us to speculate that PH domain dimerization might be a common regulatory mechanism of PH domain-containing enzymes that function on membranes.

Some receptor tyrosine kinases, such as epidermal growth factor receptor (EGFR), self-associate into high-order oligomers as part of the activation process.^45,46^ We have not studied whether dimers of the PH-TH module might form high-order oligomers on membranes, but we note that such high-order oligomers, if they exist, could further strengthen the stability of the Saraste dimer interface and contribute to the activation of Btk.

### The Saraste dimer interface and the peripheral binding site in Btk are less conserved in early stages of B-cell evolution

Neither the Saraste dimer interface nor the peripheral binding site in Btk is conserved in the PH-TH modules of Tec and Itk, the other two members of the Tec family of kinases. This raises the question of when the PIP_3_-dependent dimerization of the PH-TH module of Btk originated.

We examined the conservation of those residues that constitute the Saraste dimer interface and the peripheral binding site, comparing the sequence of human Btk with that of other mammals (105 species), birds (47 species), and fish (41 species). We found that these residues are for the most part well conserved in mammals and birds, and much less extensively conserved in fish (Figure S8). In contrast, PI3K (the enzyme responsible for generating PIP_3_) and residues in the canonical binding site of Btk are conserved in all eukaryotes. This analysis suggests that membrane binding is a more ancient function than allosteric binding for PIP_3_ in its interactions with the PH domain. The origins of the adaptive immune system, in particular B cells, are thought to be linked to the emergence of jawed fish, such as Chondrichthyes and Teleost fish^47,48^. ‘ B cells in fish have antigen-specific IgM responses similar to those in mammals, but the time required for fish B cells to generate a significant antigen-specific response is generally much longer than that required in mammals,^48^ suggesting that the signal transduction pathways that control the antigen-specific response, in which Btk plays an important regulatory role, are much less sensitive in fish than in mammals. One possibility is that the utilization of a PIP_3_-dependent dimerization mechanism for Btk activation in mammals is a consequence of natural selection for high sensitivity of the B-cell response.

### Novel strategies for developing Btk inhibitors that are selective and can inhibit the ibrutinib-resistant C481S mutant

A number of ATP-competitive Btk inhibitors have been identified in recent years.^49,50^ The most clinically tested of these is ibrutinib,^51^ which has been approved by the FDA for the treatment of mantle cell lymphoma (in 2013), chronic lymphocytic leukemia (in 2014), and a particular form of non-Hodgkin lymphoma (in 2015). Despite ibrutinib’s high efficacy in the treatment of multiple B-cell cancers, ibrutinib-resistant mutations—in particular, C481S—have emerged in a substantial fraction of all chronic lymphocytic leukemia patients treated with ibrutinib.^52^ In addition, ibrutinib has off-target effects on EGFR, ITK, and Tec-family kinases, which may result in various adverse effects in some patients.

The emerging resistance to and off-target side effects of ibrutinib have led to the active development of second-generation and more specific Btk inhibitors. Our finding that PIP_3_-dependent dimerization may be a feature of Btk activation suggests a potential new approach to the development of such inhibitors in which small molecules disrupt the dimerization of PH-TH modules. Such molecules could potentially bind either at the Saraste dimer interface or in other regions on the PH-TH module that could allosterically impair Saraste dimer formation. Inhibitors targeting the PH-TH module should act on both wild-type Btk and the C481S mutant, and should not interfere with functions of other kinases, thus addressing both of these ibrutinib-related shortcomings at the same time.

## Acknowledgments

The authors thank Konstantin Yatsenko, Thomas Weinreich, Cristian Predescu, and Tamas Szalay for helpful discussions and help with tempered binding simulations, Je-Luen Li for help with parameterization of lipids, Yibing Shan and Michael Eastwood for a critical reading of the manuscript, and Berkman Frank and Jessica McGillen for editorial assistance.

